# PBAP chromatin remodeler mediates enhancer-driven transcription in *Drosophila*

**DOI:** 10.1101/2020.04.11.036772

**Authors:** Y.V. Shidlovskii, A.V. Shaposhnikov, O.V. Bylino, D. Amendola, G. De Simone, N. Afeltra, P. Schedl, F.A. Digilio, E. Giordano

**Author notes:** Contacs: Shidlovskii Y.V., Shaposhnikov A.V., Bylino O.V., Amendola D., De Simone G., Afeltra N, Schedl P., Digilio F.A., Giordano E.

## Abstract

Chromatin remodeler SWI/SNF is an important participant of gene activation acting predominantly by opening chromatin structure on promoters and enhancers. Here we describe its novel mode of action by mediating targeted action of enhancer. We studied functions of two signature subunits of PBAP subfamily, BAP170 and SAYP, in *Drosophila*. These subunits were stably tethered onto transgene reporter carrying *hsp70* core promoter. Tethered subunits mediate transcription of reporter in a pattern prescribed by nearby enhancer in multiple loci throughout the genome, where the reporter construct was located. Both tethered SAYP and BAP170 recruit the whole PBAP complex onto reporter promoter. Studying difference between these subunits, we found that BAP170-dependent transcription is more resistant to depletion of other PBAP subunits, what may imply the principal role of BAP170 in establishing enhancer-dependent transcription.

**Author Summary:** Chromatin remodelers are key molecular machines that are responsible for local changes in chromatin structure in the nucleus. However, their functions in gene expression regulation seem to be broader. We describe the involvement of the SWI/SNF family of remodelers in establishing enhancer-promoter communication, which is apparently independent of its local remodeling activity.

Using an artificial tethering of a remodeler on a promoter, we demonstrated that promoters of a certain type become responsive to activation by a nearby enhancer only in the presence of the remodeler, while a remodeler tethering itself is insufficient for gene activation. Thus, our approach helps to uncover novel aspects of molecular interplay on regulatory elements during the gene activation process.

## Introduction

The evolutionary conserved SWI/SNF class of chromatin remodelers plays essential roles in multiple processes of cell biology, from the regulation of transcription to chromosome segregation, DNA replication, and DNA repair [1–3]. In the transcription process, SWI/SNF is important for a local increase of DNA template accessibility by physical remodeling of chromatin structure [4–6]. ATPase activity of the central subunit of the complex mediates nucleosome sliding, eviction or disassembly [7–9].

In *Drosophila*, as in multiple species, the SWI/SNF remodeler, the Brahma complex, exists in two different forms, BAP (BAF in human and SWI/SNF in yeast) and PBAP (PBAF in human and RSC in yeast) [10]. A common core complex includes Brahma (BRM, SMARCA2/4 in human and STH1/SNF2 in yeast), Moira (MOR, SMARCC1/2 in human and SWI3/RSC8 in yeast), and Snr1 (SMARCB1 in human and SNF5/ SFH1 in yeast) and can associate with the distinctive subunit OSA (ARID1A/B in human and SWI1 in yeast) to form the BAP complex or, alternatively, with Polybromo (PB, PBRM1 in human and RSC1/2/4 in yeast), BAP170 (ARID2 in human and RSC9 in yeast), and SAYP (PHF10 in human, no homolog in yeast) to produce the PBAP form [11, 12]. Analysis of mutations affecting the signature subunits has demonstrated that BAP and PBAP execute distinct and partly antagonistic functions in transcription control and development with BAP being mainly involved in cell cycle regulation and PBAP, in signal transduction cascades and differentiation [12–17]. Interestingly, the PBAP subunits have a hierarchical role in the stability of the complex. MOR is strictly required for complex core assembly, and subunits that fail to assemble into a complex are quickly degraded [12]. BAP170 is required for the stability of PB [12, 13], and the stability of BAP170, in turn, depends on SAYP [18].

To study the local functions in transcription for a factor of interest, the factor is possible to artificially tether to DNA. In yeast, two subunits of SWI/SNF have been tested in this way: LexA:SNF5 and LexA:SNF2 fusions acted as a potent transcription activator of the LexAop-carrying reporter, and activity of each fusion strongly depended on SNF2, SNF5 and SNF6 [19, 20]. Similarly, a Gal4-SAYP fusion activated a UAS-carrying reporter in a PB- and BAP170-dependent way in *Drosophila* [21]. However, the reporter was located on a plasmid in these experiments.

To better characterize the local function of the PBAP chromatin remodeler in transcription in the context of chromatin, we used a LexA:LexAop-mediated in vivo recruitment of the PBAP accessory subunits SAYP and BAP170. We found that the recruitment of the complex just upstream of a genome-integrated reporter core promoter is not sufficient to activate transcription. Conversely, its activation requires interplay between transgene-targeted PBAP and active genomic enhancers located nearby the reporter insertion site. The interplay results in activation of the target promoter and a tissue-specific expression pattern characteristic of the enhancer trapped. Both proteins efficiently recruit other PBAP subunits onto the promoter. However, an analysis of the enhancer capture effect mediated by targeted BAP170 or SAYP combined with in vivo depletion of specific PBAP subunits demonstrated that BAP170 is most likely responsible for mediating enhancer-promoter communication.

## Results

### In vivo targeting of SAYP or BAP170 to a minimal promoter mediates enhancer-dependent transcriptional activation

In a whole organism, tight gene expression control in space and time is achieved via specific interactions between enhancers and core promoters. The role of the PBAP complex in such environment has never been explored. It remains unclear how the PBAP complex functionally integrates its activity with promoter- or enhancer-bound transcription factors in the genome context.

To address these questions, we decided to use an in vivo approach to reproduce the recruitment of the PBAP complex to the promoter in transgenic flies, using both SAYP and BAP170 subunits. Specific responder and driver transgenic lines were designed using the LexA/LexAop binary system (Fig 1). In the responder transgene *LexAop:LacZ*, a *LacZ* reporter was placed under the control of a minimal *hsp70* promoter (lacking the GAF binding sites) and 8x LexA binding sites were placed upstream of the *hsp70-lacZ* fusion (Fig 1A). This construct was used to obtain 20 independent transgenic lines, which were examined for the genomic insertion sites and orientations of the constructs. Driver transgenes for expression of LexA:SAYP or LexA:BAP170 were prepared using either the ubiquitous alpha-tubulin promoter (P*tub*) or the less potent BAP170 promoter (P*BAP170*) to simulate nearly physiological levels of both subunits (Fig 1B) [16]. As a control, a transgene was constructed to ensure ubiquitous expression of the LexA repressor under the P*tub* promoter (Fig 1B). Once transgenic stocks were established for each driver construct, we verified proper expression of the constructs by immunostaining of larval tissues (S1A,B Figs) and checked whether the fusion proteins preserved their wild-type functions. For LexA:BAP170, we found that both types of driver transgenes were able to fully rescue mutant lethality of the *BAP170hfl1* allele. For LexA:SAYP, we found that the fusion binds to the same polytene chromosome sites as the endogenous SAYP protein (S2 Fig), indicating its proper recruitment to chromatin. Each *LexAop-LacZ* responder was combined with either the P*tub-LexA:SAYP* or P*tub-LexA:BAP170* transgene, and F1 third-instar larval tissues were monitored for *LacZ* expression for each cross (Fig 1C).

**Figure 1.**
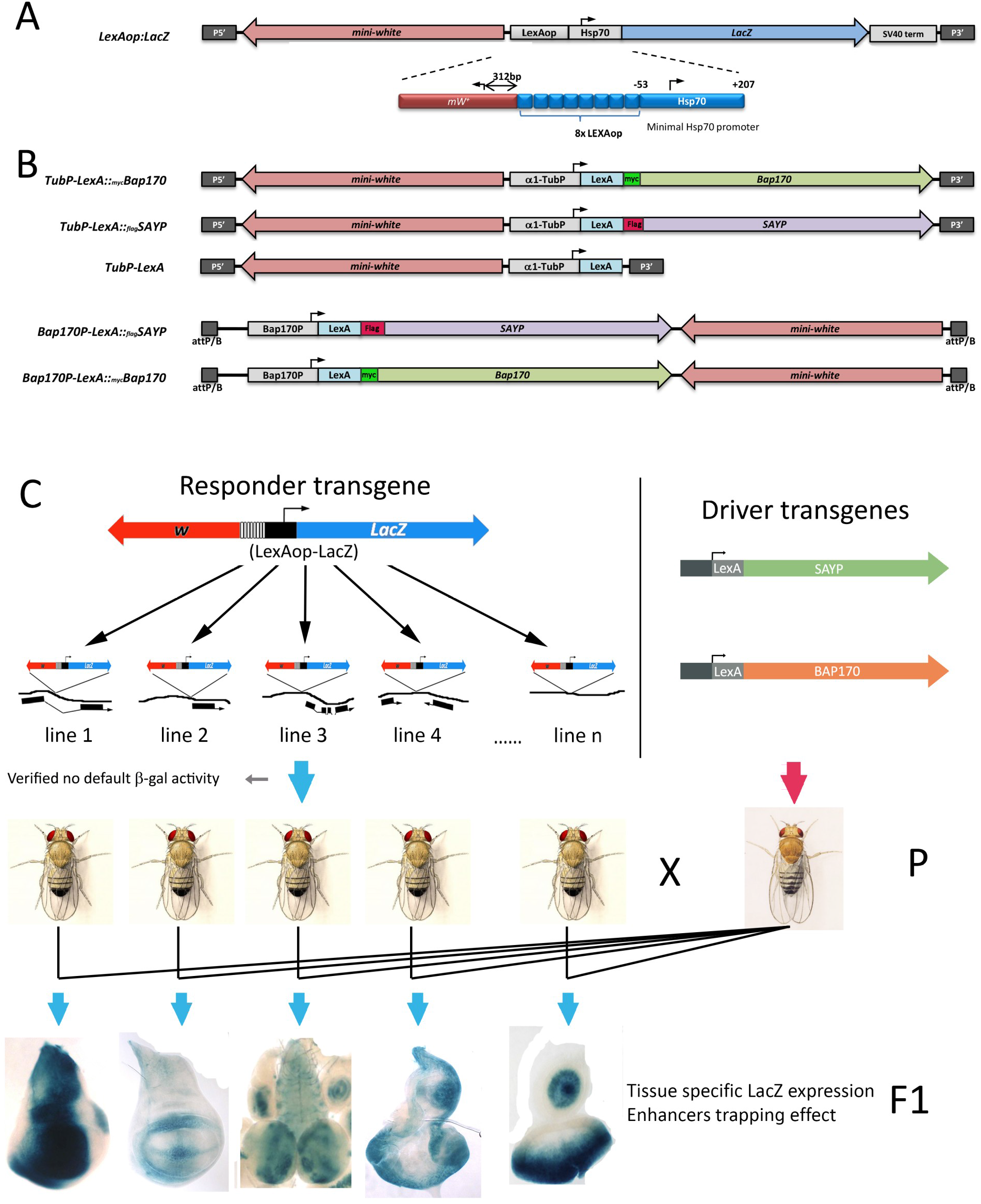
Schematic representation of the transgenes and rationale for in vivo targeting of the PBAP to a reporter promoter. (A) The P element-based *LexAop:LacZ* responder construct includes not only the dominant marker *white* gene, but also the *LacZ* reporter driven by the core promoter (−44 to +204) of the *Drosophila hsp70* gene. The region upstream of the *hsp70* promoter harbors 8x *E.coli* LexA repressor binding elements (LexAop operators). (B) Scheme of the driver transgenes designed for LexA:BAP170, LexA:SAYP, and LexA ubiquitous expression ensured by the alpha-tubulin gene promoter (P element-based constructs, top) or the promoter of the *BAP170* gene (attB/P-based construct, bottom). (C) Rationale of the procedure (see text) used to test the effect of PBAP targeting to the core promoter of the LexAop-LacZ reporter transgenes inserted in different sites of the *Drosophila* genome.

We observed that targeted SAYP or BAP170 (tSAYP or tBAP170 hereafter) acted in nearly identical manner on expression of the given *LexAop-LacZ* responder in the respective line. Surprisingly, although the LexA fusion proteins were ubiquitously expressed under the control of P*tub*(Fig S1), none of the responder lines showed corresponding ubiquitous beta-gal activity. Conversely, twelve out of the twenty *LexAop-LacZ* lines efficiently expressed the *LacZ* reporter. Yet its expression was not ubiquitous but followed a precise and reproducible tissue-specific pattern that was unique for each line (Figs 2A and S3). The *LacZ* expression patterns were reminiscent to the common enhancer trap lines with the exception that they depended on tSAYP or tBAP170. For example, the *LacZ* expression pattern of some lines clearly reproduced the expression profile of known genes, such as *Dad* (*Daughters against dpp*, [22]), *dpp* (*decapentaplegic*, [23]), *dan* (*distal antenna;* [24]) and *tara* (*taranis*, [25]) (Figs 1C and 2A). Finally, the remaining eight *LexAop-LacZ* lines showed no appreciable *LacZ* expression in third-instar larval tissues when combined with the LexA:BAP170 orLexA:SAYP drivers (not shown), demonstrating that tSAYP or tBAP170 alone are insufficient for transcriptional transgene activation once in vivo recruited nearby a promoter. In the control, LexAalone did not induce any expression of the responsive *LexAop-LacZ* transgenes (Figs 2A and S1).

**Figure 2.**
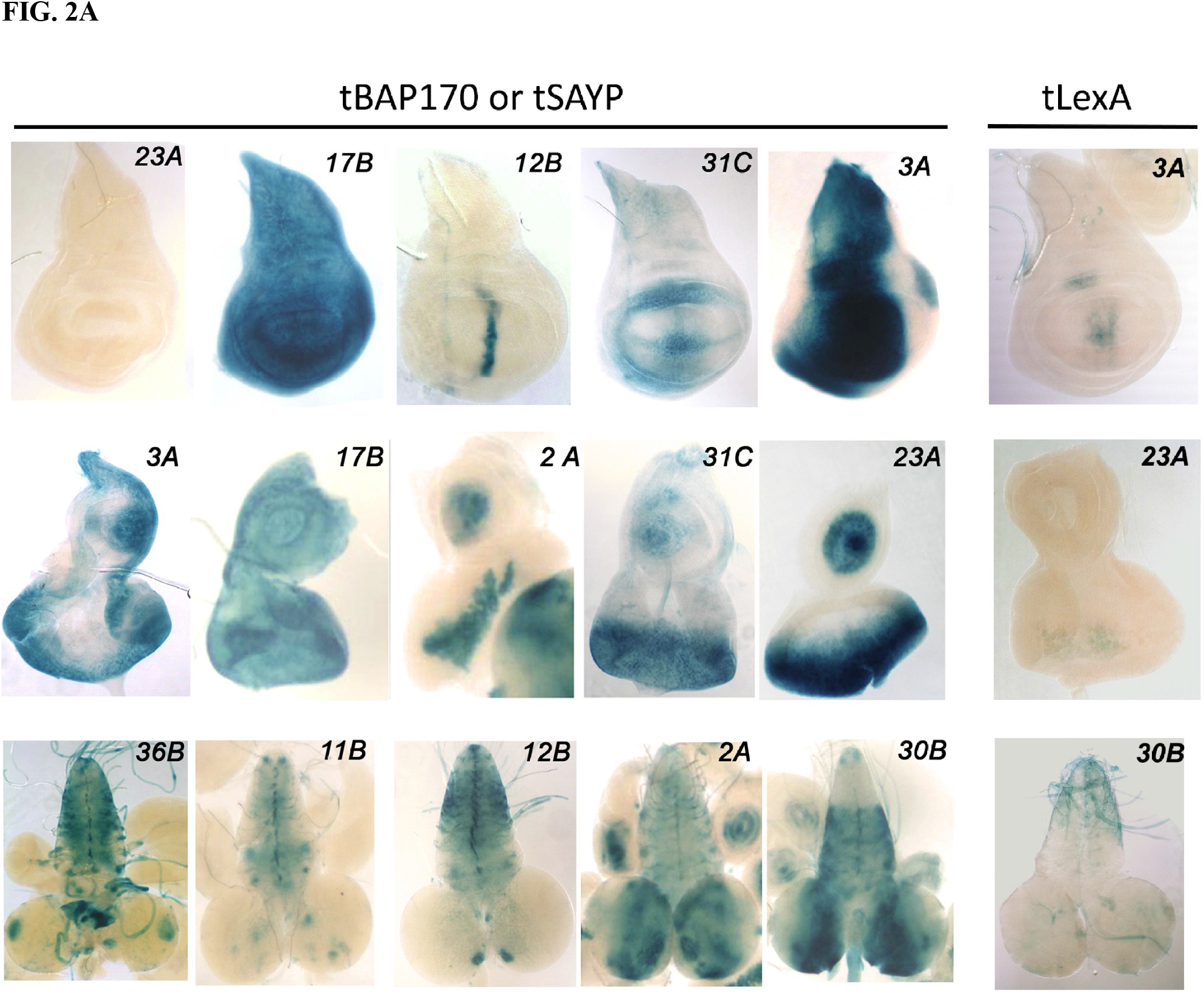
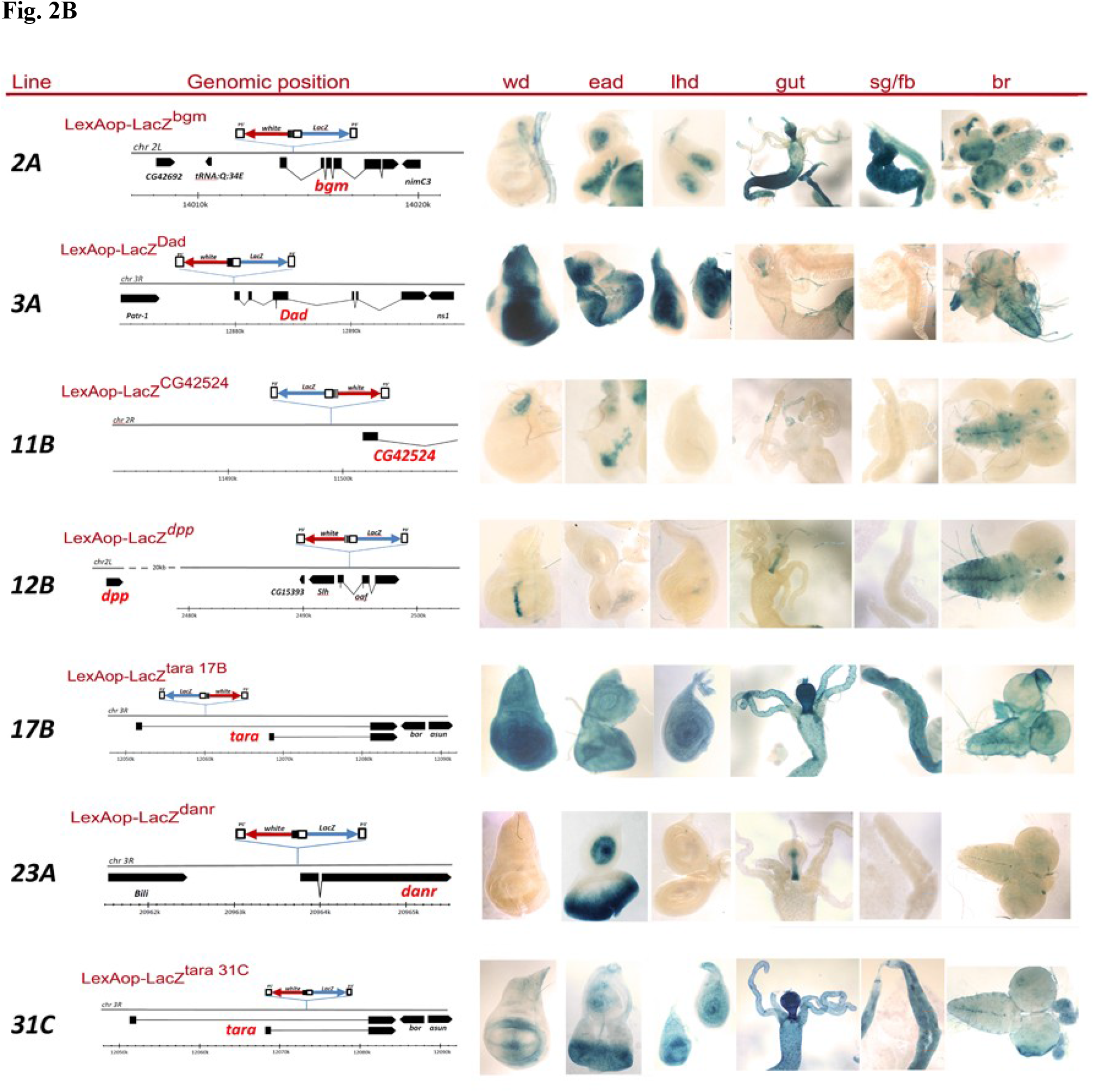
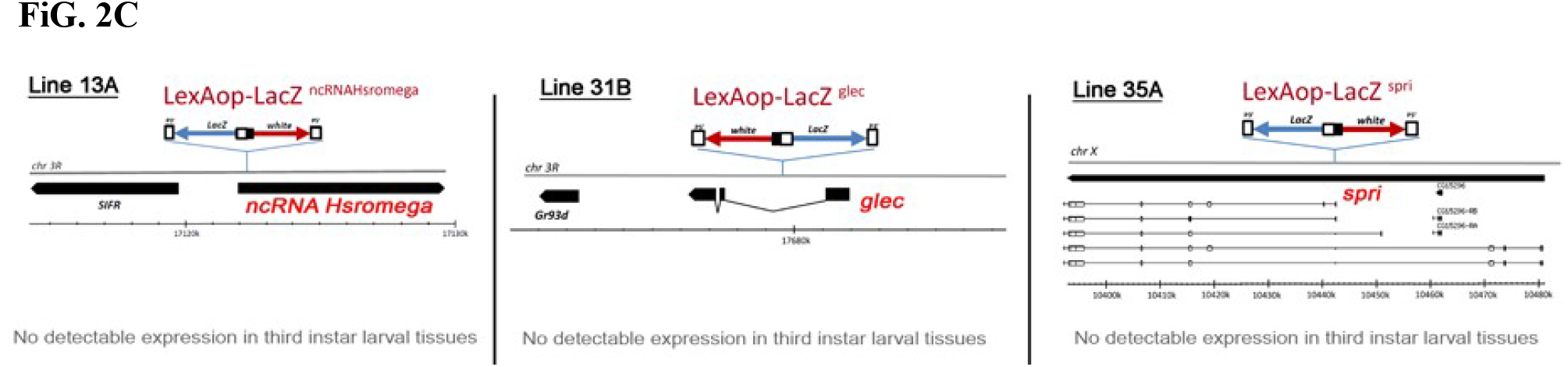
Enhancer capture by targeted BAP170/SAYP at the promoters of independent reporter transgenes. (A) A detailed view of the tissue-specific expression patterns obtained after introducing P*tub*- LexA:BAP170/SAYP to different independent lines expressing the *LexAop-LacZ* responder element in the wing discs (top), eye-antennal discs (middle), and larval brain (bottom). Note that the expression pattern was *Dad*-like in line 3A, *dpp*-like in 12B, *danr*-like in 23A, and *tara*-like in 17B. Conversely, no expression was induced by targeting the LexA protein alone (lines 3A, 23A and 30B are shown as an example) (B) Genomic positions of the integrated *LexAop-LacZ* elements in the most representative lines are shown (left) together with the corresponding P*tub*-BAP170-induced *LacZ* expression patterns in third-instar larval tissues (right). Wd, wing imaginal discs; ead, eye-antennal imaginal discs; lhd, leg-haltere discs; g, guts; sg/fb, salivary glands/fat bodies; br, larval brain. (C) Genomic positions of the *LexAop-LacZ* element in three non-responsive lines.

Because none of the responsive *LexAop-LacZ* transgenes showed significant beta-gal activitywithout the LexA fusion protein as an activator, we concluded that, once recruited to the reporter LexAop sequences, tSAYP/tBAP170 can induce “enhancer capture” by instructing the *hsp70* minimal promoter to fully respond to enhancer elements located nearby the transgene insertion site.

Once the insertion sites of these lines were determined, the enhancer-capture effect was clear (Fig 2B). For example, the reporter element was inserted close to the *Dad* gene (Figs 2B and S4) about 5 kb upstream of the mapped *Dad* enhancer [26] in *LexAop-LacZDad* or within the *oaf* gene 15kb downstream of the *dpp* disc enhancer [27] in *LexAop-LacZdpp* (Figs 2B and S4). We asked additionally if the enhancer capture effects could be a result of a global transcriptional activation induced by a potential accumulation of PBAP remodelers caused by overexpression of their key signature components, such as SAYP and BAP170. Therefore, we checked if wild type SAYP or BAP170 could induce activation of the *LexAop-LacZ* responders similarly to LexA:SAYP or LexA:BAP170. We found that none of the responsive *LexAop-LacZ* reporters was activated in cells overexpressing the wild-type BAP170 or SAYP (S5 Fig), demonstrating that any eventual increasein untargeted PBAP was not responsible for the enhancer-dependent activation of the *LexAop* reporters.

In summary, these data indicate that the tSAYP/tBAP170-induced enhancer responsiveness has characteristics similar to the normal enhancer-promoter communication, (i) being achievable with both upstream and downstream enhancers and (ii) acting at a long distance.

### Promoter-bound PBAP subunit functions as a tethering factor for the *hsp70* minimal promoter

The standard *hsp70* minimal promoter used in the *LexAop-LacZ* transgene (sequence from −44 to +204) has intrinsic responsiveness to several enhancers when they are located in its 5’ proximity[28–30], but responds less efficiently when they are at a greater distance [31]. Conversely, many other promoters used in transgenic analyses of enhancer/promoter interactions, such as the *mini-white* and P-transposase promoters, have the ability to interact at a distance as well, being naturally provided with promoter-tethering elements [29, 32, 33]. Accordingly, all the enhancers identified by the tBAP170/tSAYP-inducible *LexAop-LacZ* responders are fully compatible with the *hsp70* minimal promoter, but, in the absence of tBAP170/tSAYP, their interactions are presumably prevented by the lack of a tethering mechanism. This is evident in the *LexAop-LacZDad* and *LexAop-LacZdanr* lines, where the *Dad* and the *danr* enhancers cannot interact with the *hsp70* promoter without tBAP170/tSAYP, but can spontaneously activate the *mini*-*white* or P-transposase promoter, as demonstrated by the *Dad*-like or *danr*-like expression pattern of the transgene *mini-white* marker in the wing and eye discs (S6 Fig).

Therefore, if tBAP170 functions as a simple tethering factor between the minimal *hsp70* promoter and distant enhancers, we expected that relocating enhancers closer to the promoter could bypass tBAP170 requirement and restore the regular enhancer-promoter interaction. Using φC31 site168 specific recombination, a series of transgenes was integrated in the same attP2 site on the third chromosome. In the transgenes, the isolated *Dad* or *dpp* enhancers were placed either just upstream of the *LexAop-hsp70-LacZ* cassettes or, as a control, downstream of the *LacZ* reporter about 4 kb from the *hsp70* promoter (Fig 3). The *Dad* and *dpp* enhancers were taken because their positions in the genome were known [26, 28]. As expected, the distant *Dad* and *dpp* enhancers activated expression of the *LacZ* reporter only in the presence of tBAP170, as already observed with the *LexAop-LacZ* reporters. Conversely, the enhancers located in close proximity of the reporter strongly activated its expression independently of tBAP170, whose presence had no evident effect on *LacZ* expression. Therefore, tBAP170 mediates the interaction between distant enhancers and the *hsp70* minimal promoter.

**Figure 3.**
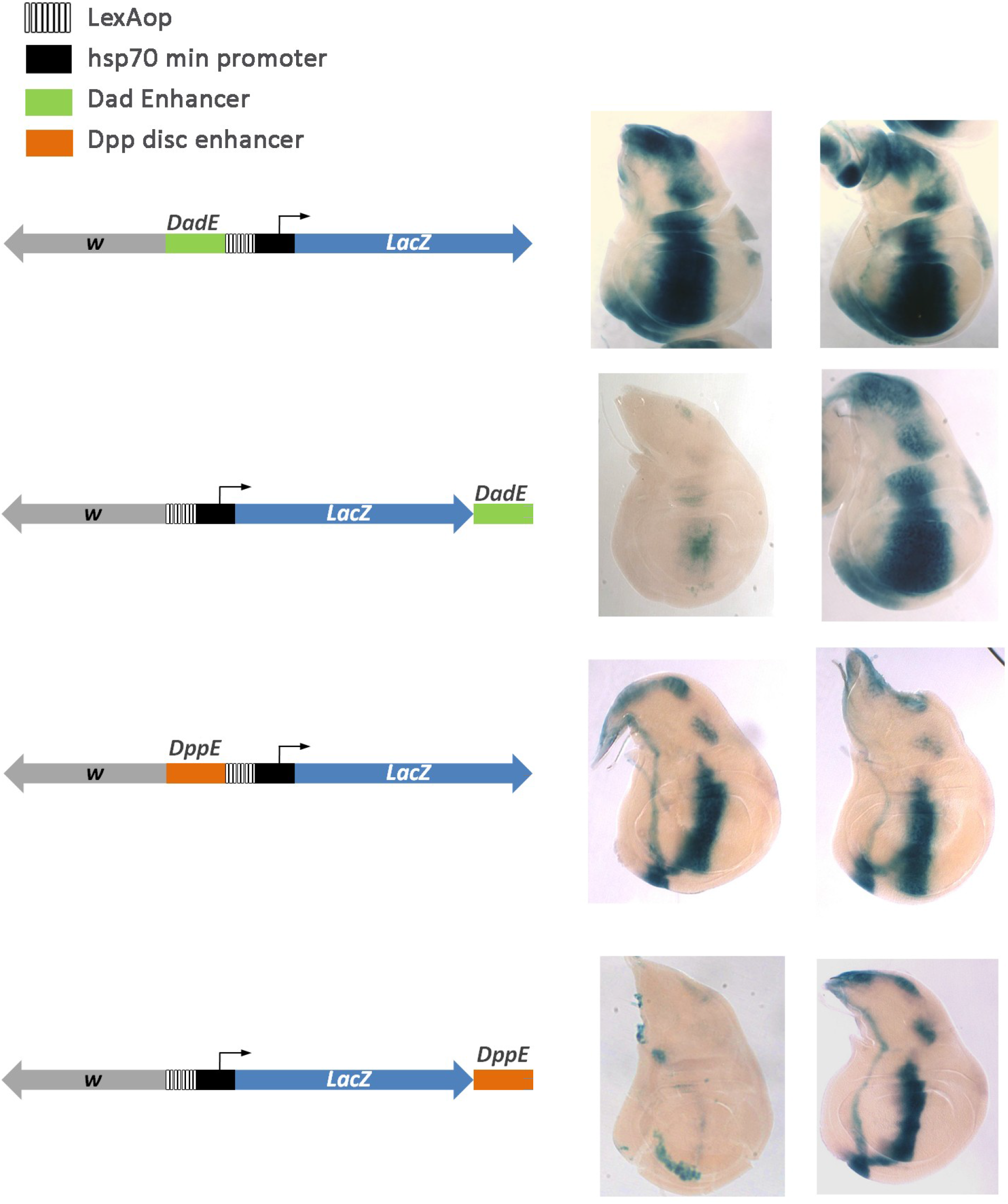
tBAP170 functions as tethering factor between the promoter and distant enhancers. Beta-Gal activity in transgenic lines with the *LexAop-hsp70-LacZ* constructs carrying the *Dad* (top) or *dpp* (bottom) enhancers. The *hsp70* core promoter can respond to both enhancers in the absence of tBAP170 when the enhancers are in close 5’ proximity (panels 1 and 3, -), but not when they are at a distance (panels 2 and 4, -). Conversely, the distant enhancers are capable of activating of the *hsp70* core promoter in the presence of tBAP170 (panel 2 and 4, +). From top to bottom, the genotypes are: *JFL-attB [DadEnh-lexAop-hsp70-LacZ], JFL-attB [lexAop-hsp70-LacZ-DadEnh], JFL-attB [dppEnh-lexAop-hsp70-LacZ], and JFL-attB [lexAop-hsp70-LacZ-dppEnh]*. The absence (-) or presence (+) of the P*tub*-LexA:BAP170 driver is indicated.

### Tethered PBAP subunit recruits the whole complex onto the *hsp70* promoter

To gain insight into the molecular mechanism of action of the tethered subunits, the transgenic locus was analyzed in ChIP experiments. We used the flies that carried the *LexAop-LacZDad* responder and P*tub*- LexA:BAP170 driver. Flies carrying P*tub*-LexA:SAYP had strongly reduced the viability, and we consequently could not collect enough material for a ChIP experiment.

The LexA fusion was efficiently recruited onto the transgenic *hsp70* promoter (Fig 4A). At the same time, a significant peak of LexA was detected on the endogenous *Dad* enhancer. The findings possibly reflect the formation of spatial enhancer-promoter contacts, leading to LexA cross linking with a distal enhancer.

**Figure 4.**
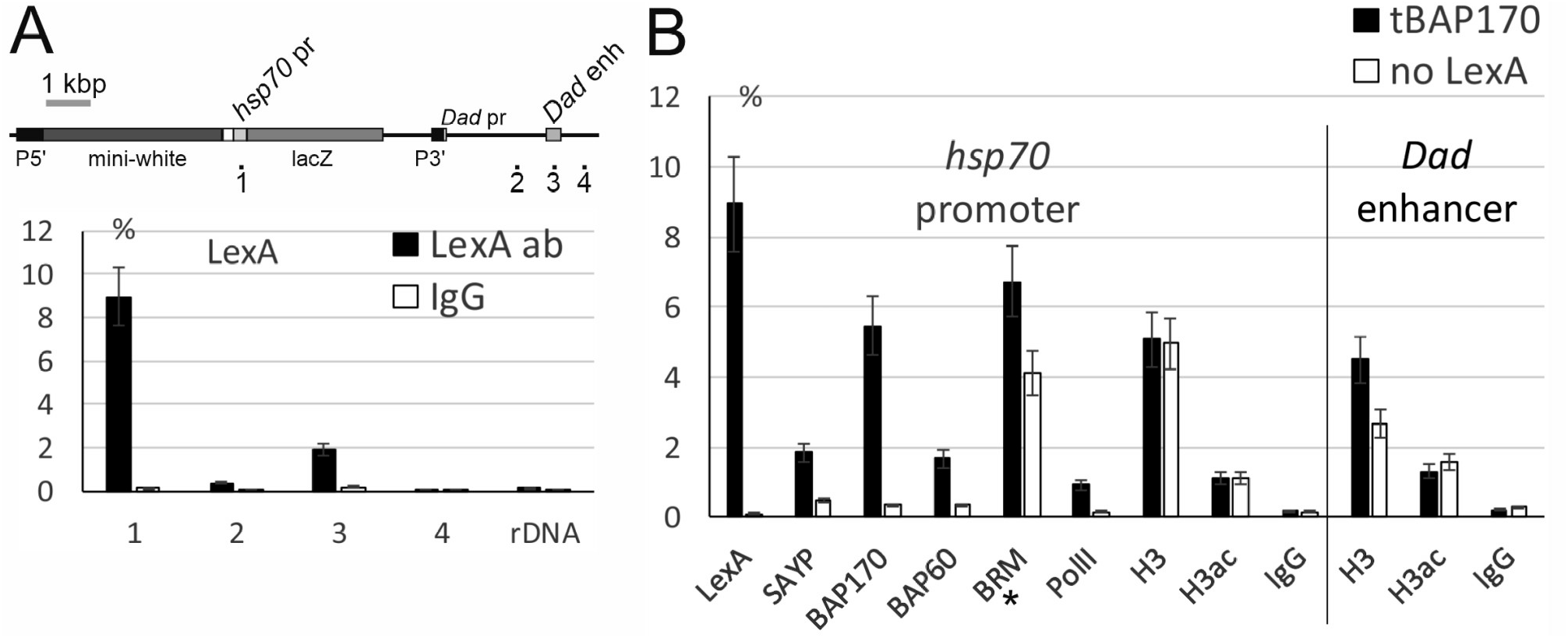
ChIP analysis of the *lexAop-LacZDad* responder line expressing the LexA:BAP170 fusion. (A) ChIP profile of LexA:BAP170 fusion binding along the transgenic *Dad* locus (a scheme is given at the top). Loci 1-4 indicate the positions in the transgenic *Dad* locus. IgG (pre-immune immunoglobulin) and rDNA (ribosomal 28S RNA gene) were used as negative controls. (B) Recruitment of different proteins onto the *hsp70* reporter promoter and *Dad* enhancer. The LexA fusion and subunits of the PBAP complex are indicated at the bottom. PolII, RNA polymerase II; H3, histone H3; H3ac, acetylated H3. no LexA, control flies, which did not express the LexA fusion. (*) The BRM level increased tenfold. The level of a factor is shown as a percentage of Input.

We have checked whether other PBAP subunits were present on the *hsp70* promoter. Endogenous SAYP and two core PBAP subunits, BAP60 and central ATPase BRM, were efficiently recruited by tBAP170 (Fig 4B). An increase in PolII corresponding to increased gene activity was also detected. Thus, the tethered subunits recruited the whole complex, eventually leading to PolII recruitment and gene activation.

Finally, we checked whether the recruited PBAP remodeler affected the chromatin state on the *hsp70* promoter and spatially close *Dad* enhancer. Despite the presence of the PBAP core, the level of histone H3 was not decreased on the active enhancer and promoter (Fig 4B). An increase in histone H3 acetylation was similarly not detected on both elements. Thus, the tethered chromatin remodeler did not create a nucleosome-free region or stimulate the imposition of an epigenetic mark of active chromatin on the *hsp70* promoter and *Dad* enhancer. The remodeling activity of the complex seems to be dispensable for mediating enhancer-dependent transcription in this case.

### tSAYP-dependent reporter transcription crucially depends on the PBAP subunits

Because recruitment of the whole PBAP complex was shown in the ChIP experiments, we checked the functional importance of other PBAP subunits in tSAYP and tBAP170-mediated transcription. The genetic system created allowed us to use the wide collection of *Drosophila* GAL4/UAS-based tools to manipulate the genetic background and make functional tests.

To check if a component the PBAP complex is necessary for tSAYP-mediated enhancer capture, we prepared a stock that contained the P*BAP170-LexA:SAYP*(*attP2*) driver and the *LexAop-LacZDad* reporter recombined on the third chromosome together with the engrailed-GAL4 (en-GAL4) and UAS-GFP elements recombined on the second chromosome. The stock continuously expressed the *LacZ* reporter according to the tSAYP-induced Dad-like pattern and was used to knock down expression of key components of the PBAP complex in the posterior compartment of the imaginal discs using the GAL4-dependent UAS-RNAi lines (S1 Table). The en-GAL4 driver was chosen among several GAL4 drivers with *Dad* overlapping expression because it was the only one that ensured a regular growth of the third-instar imaginal discs under an RNAi regimen and was strong enough to cause an RNAi phenotype (see below). A shown in Fig 5, the overlap between the engrailed and the Dad-like expression patterns corresponds to a large stripe of posterior cells adjacent to the A/P boundary.

**Figure 5.**
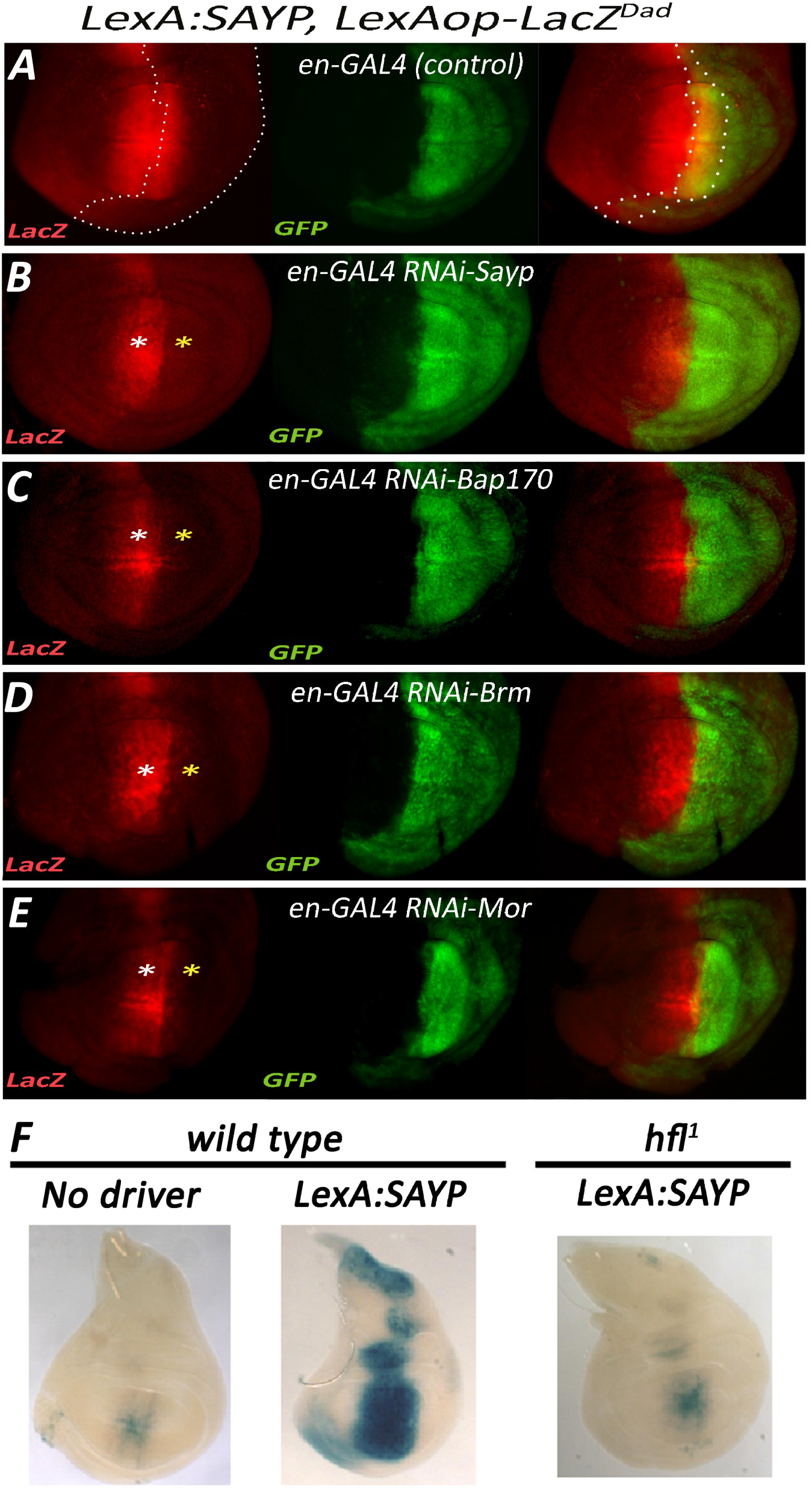
Enhancer-dependent transcriptional activation mediated by SAYP requires BAP170 and key components of the PBAP complex core. Fluorescence imaging of beta-gal (red) and GFP (green) expression in wing discs from larvae of the genotype *en-GAL4, UAS-GFP; PBAP170-LexA:SAYP, LexAop-LacZDad* transgenes alone (A control) or in combination with RNAi lines for SAYP (B), BAP170 (C), Brm (D), and Mor (E). (A) Control wing disc from *en-Gal4, UAS-GFP; PBAP170-LexA:SAYP, LexAop-LacZDad* without RNAi. The Dad like expression pattern of the SAYP-induced *LexAop-LacZDad* transgene overlaps posterior expression of en-GAL4 in a GFP-positive posterior row of cells flanking the A/P axis (cells included in the dotted line). White asterisks indicate the regions with normal *LexAop-LacZDad* transgene expression (outside the induced RNAi region), and yellow asterisks indicate the regions with expression of RNAi lines and altered target *LexAop-LacZDad* transgene activation. Genotypes: (B) *en-GAL4, UAS-GFP; PBAP170-LexA:SAYP, LexAop-LacZDad/UAS-RNAi-SAYP [Vdrc105946]*, (C) *en-GAL4,UAS-GFP; PBAP170-LexA:SAYP, LexAop-LacZDad/UAS-RNAi-BAP170 [Vdrc34582]*, (D) *en-GAL4,UAS-GFP; PBAP170-LexA:SAYP, LexAop-LacZDad/UAS-RNAi-Brm [Vdrc37721]*, (E) *en-GAL4, UAS-GFP; PBAP170-LexA:SAYP, LexAop-LacZDad/UAS-RNAi-Mor [Vdrc6969]*. (F) Beta-gal activity in wing discs from larvae with the *LexAop-LacZDad* responder alone (left) or combined with P*BAP170-LexA:SAYP* in the wild type (center) or the *BAP170* null mutant background *hlf1/hlf1* (right).

Depletion of the PBAP complex core subunit MOR or ATPase BRM was found to abolish expression of *LacZ*, implying that their presence on the reporter promoter is necessary for enhancer dependent transcriptional activation by tSAYP. Surprisingly, BAP170 depletion similarly abolished tSAYP-mediated reporter transcription, demonstrating that tSAYP alone is insufficient without the endogenous BAP170. This was confirmed by the finding that tSAYP failed to activate the transgene reporter in the null BAP170 *hfl1* mutant background (Fig 5F).

All of the effects described were transgene specific and were not a consequence of potential Dad transcription defects or functional loss of the *Dad* enhancer caused by PBAP subunit depletion. In fact, both *Dad* enhancer activity and *Dad* transcription are independent of PBAP (S7 Fig), as already described for the other *Dpp* target *spalt* [17]. In conclusion, our data indicate that the PBAP subunits tested (MOR, BRM, and BAP170) are necessary for an enhancer to mediate the induction of expression from the transgene promoter.

### tBAP170-driven transcriptional activation is resistant to depletion of PBAP core subunits

To check if tBAP170-mediated enhancer capture also requires components of the PBAP complex, as is the case with tSAYP, we prepared a stock that was similar to that described for tSAYP except that the driver expressed LexA:BAP170. The stock carried the P*BAP170-LexA:BAP170(attP2*) driver and the *LexAop-LacZDad* reporter recombined on the third chromosome together with the en-GAL4 and UAS-GFP elements recombined on the second chromosome. Both tSAYP and tBAP170 drivers were integrated in the same genomic AttP2 docking site favorable for robust expression with no position effect [23, 34, 35]. Differently from tSAYP, RNAi-mediated depletion of either BRM or MOR did not cause loss of *LacZ* expression (Fig 6). Surprisingly, even the SAYP knock down showed no effect on expression of the transgene, which is normally active in posterior cells of the wing disc (Fig4). In summary, these data indicate that tBAP170 could be sufficient to trigger the enhancer capture effect independently of the core PBAP components and SAYP and that, within the PBAP complex, BAP170 may represent a subunit with tethering activity required to confer enhancer promoter interactions.

**Figure 6.**
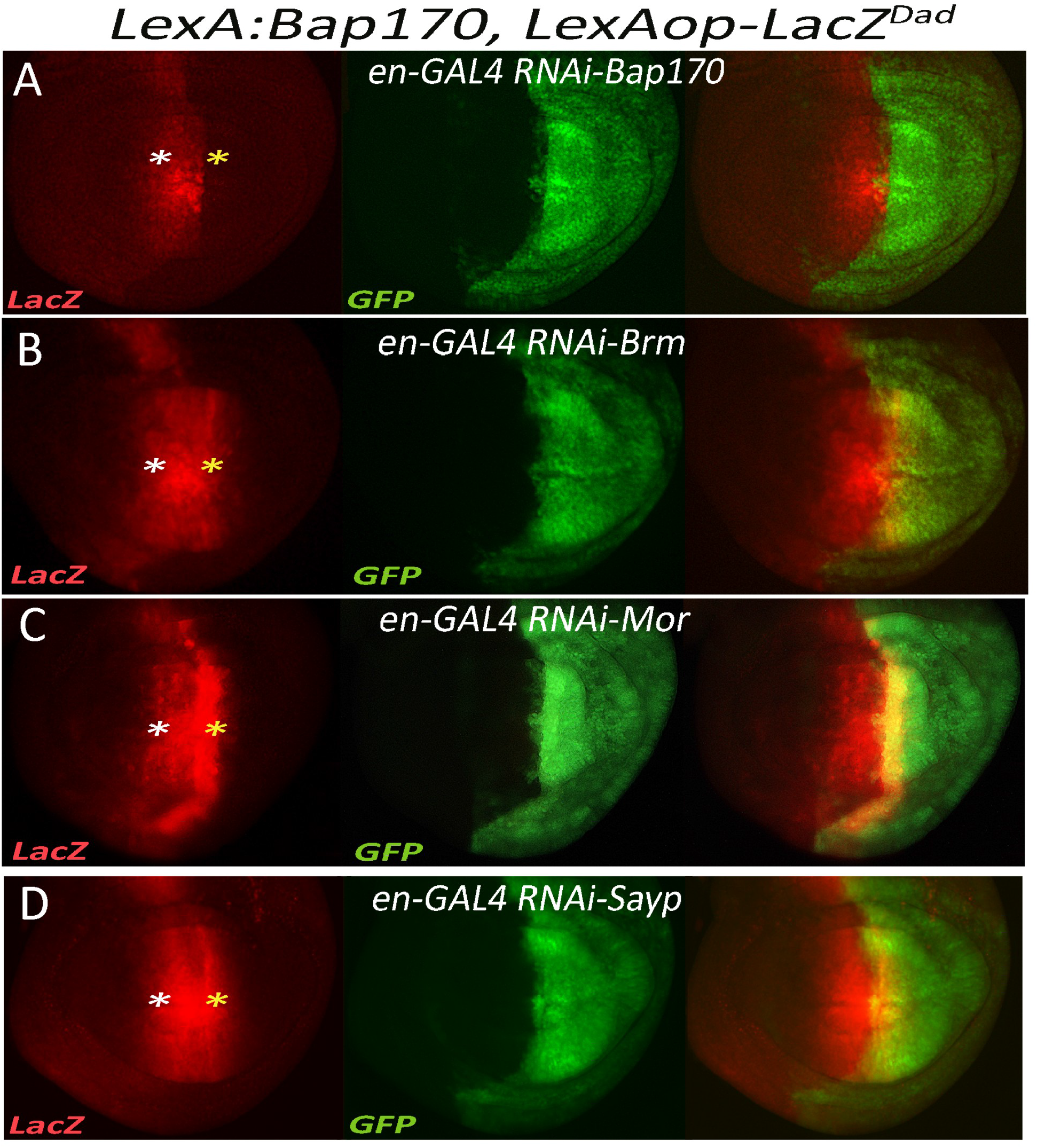
Targeted BAP170 can capture the enhancer independently by SAYP and the PBAP complex core. Fluorescence imaging of beta-gal (red) and GFP (green) expression in the wing discs from larvae of the genotype *en-GAL4, UAS-GFP; BAP170P-LexA:BAP170, LexAop-LacZDad* transgenes in combination with RNAi lines for BAP170 (A), Brm (B), Mor (C), and SAYP (D). White asterisks indicate the regions with normal *LexAop-LacZDad* transgene expression (outside the region of RNAi induction), and yellow asterisks indicate the regions that overlap the RNAi line activation. Genotypes: (A) *en-GAL4, UAS-GFP; PBAP170-LexA:SAYP, LexAop-LacZDad/UAS-RNAi-BAP170 [Vdrc34582]*, (B) *en-GAL4, UAS-GFP; PBAP170-LexA:SAYP, LexAop-LacZDad/UAS-RNAi-Brm [Vdrc37721], (C)en-GAL4, UAS-GFP; PBAP170-LexA:SAYP, LexAop-LacZDad/UAS-RNAi-Mor [Vdrc6969]*, (D) *en-GAL4, UAS-GFP; PBAP170-LexA:SAYP, LexAop-LacZDad/UAS-RNAi-SAYP [Vdrc105946]*.

## Discussion

Here we showed that artificial tethering of signature subunits of the PBAP chromatin remodeler onto the core *hsp70* promoter induces expression of a downstream reporter in a way dictated by the nearby enhancer. Enhancer trapping by the promoter occurs only when a PBAP subunit is tethered on the promoter, suggesting a crucial role in enhancer-promotor communication for the remodeler.

It is known that SWI/SNF targeting to chromatin could be executed by a core subunit [36, 37]. However, the signature subunits seem to have greater importance for correct recruitment of the complex [38, 39]. Thus, our system reproduces the endogenous mechanisms of recruitment in a correct way. Indeed, the whole complex was detected on the *hsp70* promoter in ChIP analysis.

The SWI/SNF remodeler is localized on promoters throughout the genome and plays an important role in the establishment of a specific nucleosome pattern on a promoter [40]. Enhancer shown even a greater requirement for SWI/SNF [37, 38, 41–48]. SWI/S NF similarly establishes the nucleosome landscape on enhancers [44, 49], and its participation in local histone acetylation has also been described [42]. Interestingly, in mammals, the PBAF subfamily shows preferable localization on promoters and BAF, on enhancers [36].

Our enhancer trapping system showed promoter specificity: the minimal *hsp70* promoter required PBAP to be activated by an enhancer, while the endogenous P-element and *white* promoters did not. Thus, recruited PBAP can confer an enhancer specificity on at least a certain subset of promoters. Promoter specificity of the kind is usually determined by various core promoter motifs, promotor proximal elements, and some epigenetic signals [50, 51]. The elements are recognized by a number of both DNA-binding and accessory proteins, among which architectural factors are of particular importance, mediating long-range interactions [52–54]. Our data indicate that PBAP is a component of this complicated system, which is necessary for establishing specific enhancer-promotor communication.

Several hypotheses are possible to advance to account for the role of tethered PBAP in facilitating the enhancer-promotor contacts in our system. An evident suggestion is that locally accessible chromatin forms on the promoter or spatially adjacent enhancer to facilitate recruitment of the transcription apparatus. However, we did not detect a decrease in nucleosome density or an increase in histone acetylation on the promoter and corresponding enhancer. Previous investigations have shown that recruitment of the remodeler to a definite site is not always accompanied by local chromatin remodeling [55, 56]. Moreover, enzymatic activity of the complex is not always important for its function: Brahma regulates a half of its target genes in *Drosophila* through a mechanism that does not require ATPase activity [57].

In this case, a structural role is possible to assume for the remodeler. That is, the promoter-bound SWI/SNF complex as a transcriptional coactivator interacts with enhancer-bound activators or some chromatin architectural factors and thus facilitates promoter-enhancer communication. Indeed, the SAYP subunit has previously been shown to interact with DNA-specific transcription activators [58, 59]. Interestingly, SAYP has been observed to act as a potent activator of a reporter in previous experiments, but the reporter was in a plasmid [21]. Our data imply that, in the genomic context, some factor present on promoters impedes SAYP-driven transcription. This repression can be revealed by some enhancer-bound activity.

In addition, there is another intriguing possibility of a direct role of ATPase activity of the complex in enhancer-promoter communication. Besides the model where the base of a chromatin loop directly forms between the enhancer and the promoter via protein-protein interactions, there is a possibility that a loop arises via its progressive extrusion [60]. The SWI/SNF remodeler acts as a directional DNA translocase [61] and has an intrinsic property to generate DNA loops on nucleosomes [62, 63]. Loops generated in vitro are of different size, up to 1200 bp [64]. In vitro, the purified SWI/SNF complex produces loops on a polynucleosome matrix in an ATP-dependent manner, and one complex is enough to remodel a series of nucleosomes. As the complex has at least two sites to interact with different DNA sequences, it has a property to bring distantly located sites into proximity [65]. In vivo Brg1 has been found to be crucial for the formation of a loop between a regulatory element and a promoter in the *alpha*- and *beta-globin* gene loci [66, 67]. At the same time, Brg1 is not involved in the formation of chromatin loops in several other loci, suggesting locus specficity for this putative mechanism. The model of SWI/SNF-mediated loop extrusion in vivo still requires further confirmation.

Apparently, our knockdown experiments challenged the necessity of the whole complex for enhancer-promotor communication: BAP170 seems to drive enhancer-dependent activation independently of other subunits of the complex, while SAYP lacks this capability. However, this fact could be attributed to different affinities of these subunits for the core complex. Indeed, SAYP seems to be an optional subunit of PBAP because it is underrepresented in Brahma complex samples [18] and its profile only partly overlaps the profiles of other Brahma subunits [21]. BAP170 as an integral component of PBAP could recruit the residual amounts of subunits after their knockdown more efficiently and could easier overcome their depletion.

Still, there is a possibility that BAP170 is directly involved in enhancer-promoter communication, utilizing its internal properties or interacting with some architectural chromatin factors to play this role. Interestingly, the SNF6 subunit of the remodeler in yeast can also support transcription of the reporter independently of other subunits of SWI/SNF when tethered onto the promoter via LexA [68]. However, this subunit is yeast specific, and it is unclear if BAP170 and SNF6 act in the same way in transcription activation. The contributions of different subunits into enhancer-promotor communication requires further investigation.

The system described here allowed us to reveal novel aspects of the role that the remodeler plays in enhancer-dependent transcription. Recent data point to a localization of the remodeler on chromatin boundaries and its role in the formation of the global chromatin structure [15, 69–71]. Our approach could be useful for studying the remodeler function on these elements to finally elucidate the full spectrum of roles that the SWI/SNF complex and its individual subunits play in gene expression regulation.

## Materials and Methods

### Plasmid preparation

The LexAop-LacZ plasmid was prepared by cloning the BglII-LexAop-*hsp70*-HindIII(filled) fragment of pJFRC18-8XLexAop2-mCD8:GFP [35] (Addgene plasmid #26225) into the BamHI/EcoRI(filled) restriction sites of the transformation vector pCasper-AUG-beta-gal [72]. The LexAop-*hsp70* cassette consists of 8x (22-bp) LexA operators followed by the *hsp70* minimal core promoter from - 45 to +207, thus lacking the upstream GAGA binding sites.

For the LexA:myc:BAP170 in-frame cassette, the NLS:LEXA (1-214aa) fragment from the pBPnlsLexA::GADflUw plasmid [35] (Addgene plasmid #26232) was cloned in frame in a myc-BAP170 (2-1688 aa) cassette. For the LexA:3xFLAG:SAYP in-frame cassette, the NLS:LexA (1-214aa) fragment from the pBPnlsLexA::GADflUw plasmid was cloned in frame in a 3xFLAG:SAYP (2-1843 aa) cassette. The P element-based transformation vectors P*tub*-LexA:myc:BAP170 and P*tub*-LexA:3xFLAG:SAYP were prepared by inserting the LexA:m yc:BAP170 or LexA:3xFLAG:SAYP cassettes in the pOP-118 vector, which contains the ubiquitous tubulin-1α gene promoter. The AttB based BAP170 promoter-driven transgenes P*BAP170*-LexA:myc:BAP170 and P*BAP170*- LexA:3xFLAG:SAYP were prepared by inserting the LexA:myc:BAP170 and LexA:3xFLAG:SAYP cassettes in the P*BAP170*-AttB vector. The P*BAP170*- AttB vector was prepared by inserting a PCR fragment containing the BAP170 transcriptional regulatory sequences (−373/+135) [16] in the pJFRC18-8XLexAop2-mCD8:GFP-derived plasmid JFR-AttB. All cloning steps, maps, and plasmids are available upon request.

Full-length BAP170 cDNA clones have been previously described [16]. The full-length SAYP cDNA (LD10526) was purchased from the *Drosophila* Genome Resource Center.

Constructs for enhancer proximity tests *JFL-attB [DadEnh-lexAop-hsp70-LacZ], JFL-attB [lexAop-hsp70-LacZ-DadEnh], JFL-attB [dppEnh-lexAop-hsp70-LacZ]*, and *JFL-attB [lexAop-hsp70-LacZ-dppEnh]* were prepared by cloning PCR fragments of the *Dad* and *dpp* enhancers in the JFL-attB vector. The JFL-attB vector has been prepared in our lab by modifying pJFRC18-8XLexAop2-mCD8::GFP as follow: (i) the NdeI/XbaI fragment corresponding to the mini-*white* cassette was recovered from pBPnlsLexA::GADflUw, filled, and cloned into the EcoRV site of pBluescript to obtain the white-PBS; (ii) the *white* cassette from pJFRC18-8XLexAop2-mCD8::GFP was removed by EcoRV/HindIII digestion and *mini-white* was reintroduced in the opposite orientation as a HindIII/BamHI(filled) fragment from white-PBS to obtain the JFK plasmid; (iii) the GFP cassette was then removed from JFK by BglII/XbaI digestion, and the *LacZ* gene was introduced as a BamHI/XbaI fragment of pCasper-AUG-beta-gal to obtain the final JFL-attB plasmid. Primers for cloning the *Dad* and *dpp* enhancers in JFL are described in Supplement.

### Beta-gal staining, in situ hybridization, and immunofluorescence

Detection of beta-gal activity for LacZ reporters was carried out according to standard protocols. Images were captured using either a Leica MZ stereomicroscope or a Reichert-Jung Polyvar microscope using incident fiber optic lights. In situ hybridizations were carried out as described in [73], with a DIG-RNA probe prepared using pBluescript-cloned PCR-amplified fragments corresponding to the *white* exon. Indirect immunofluorescences were carried out according to standard protocols.

### Transgenic line preparation

P element-based transgenic lines were generated by injections into *w1118* embryos as previously described [74], using the transposase activity provided by the helper plasmid Turbo Δ2–3 [75].Chromosomal insertion sites of the LexAop-LacZ lines were mapped by inverse PCR with P-element end primers. The PCR fragments were then sequenced, and the insertion sites mapped via BLAST searches at Flybase. In the case of attB-basewd vectors, transgenic lines were prepared using phiC31 attB/attP site-specific recombination by injections into embryos of the genotype *y,w,P{y[+t7.7]=nos-phiC31\int.NLS}X; P{y[+t7.7]=CaryP}attP2*.

### Drosophila strains

The following GAL4 and UAS lines were used: *en-GAL4, UAS-GFP; Ms1096-GAL4; UAS378 GFP; UAS-RNAi-SAYP [Vdrc105946]; UAS-RNAi-BAP170 [Vdrc34582]; UAS-RNAi-Brm [Vdrc37721]; UAS-RNAi-Mor [Vdrc6969]*. The RNAi lines were chosen on the basis of their ability to induce clear defects with tubulin-GAL4 and en-GAL4 drivers, as well with other GAL4 lines in the literature (see Supplementary Table 1). The *P{lacW}Dadj1E4* was used as a *Dad*-LacZ enhancer trap.

### Antibodies

Primary antibodies against beta-gal (Promega), LexA (Millipore), FLAG (Sigma), myc, GFP (Sigma), PolII CTD, histone H3 (Abcam), H3ac (ab47915, Abcam), SAYP [76], BAP170 [18], and BAP60 [77] were used. Antibodies against fragment 652-785 of the RA form of BRM were raised inrabbits and affinity purified (S8 Fig). In immunofluorescence experiments, secondary antibodies antimouse Alexa Fluor 568 and anti-rabbit Alexa Fluor 488 (Thermofisher) were used.

### Chromatin immunoprecipitation

Whole larvae were taken for analysis as described in [78]. Briefly, the larvae were homogenized in a NU-1 buffer (15 mM HEPES-KOH, pH 7.6, 10 mM KCl, 5 mM MgCl2, 0.1 mM EDTA, 0.5 mM EGTA, 0.35 M sucrose, 1 mM DTT) supplemented with 0.5 % formaldehyde. The nuclear suspension was filtered through 50 μm Filcon, the filtrate was incubated for a total of 10 min at RT, quenched with an equimolar amount of glycine for 5 min at RT, and nuclei were centrifuged 1000g for 1 min. The pellet was washed 2 times with PBS and resuspended in 300 μl of a Sonication buffer (50 mM HEPES-KOH, pH 7.9, 140 mM NaCl, 1 mM EDTA, 1% Triton X-100, 0.1% deoxycholate Na, 0.1% SDS); the suspension was sonicated and centrifuged. DNA (3-10 μg) was taken for a ChIP reaction; immunoprecipitation was performed as described in [79]. Primers for qPCR are given in Supplement.

## Acknowledgements

We thank Daniela Cavaliere and Orsolina Petillo for their technical support and Dr. Mariarosaria Aletta (CNR) for bibliographic support. We are grateful to the Bloomington stock Center for *Drosophila* strains.

This study was performed using DM6 Leica microscopes at IBBR and the infrastructure of the Center for Collective Use ‘Biology of the Living Cell and Drug Biomedical Nanotransporters’ of the Institute of Gene Biology, Russian Academy of Sciences. We also thank the Center for Precision Genome Editing and Genetic Technologies for Biomedicine (Institute of Gene Biology) for the equipment and genome-editing technology.

