## Supplemental Data for "PBAP chromatin remodeler mediates enhancer-driven transcription in *Drosophila*"

### Supplementary information

#### Supplementary Figures

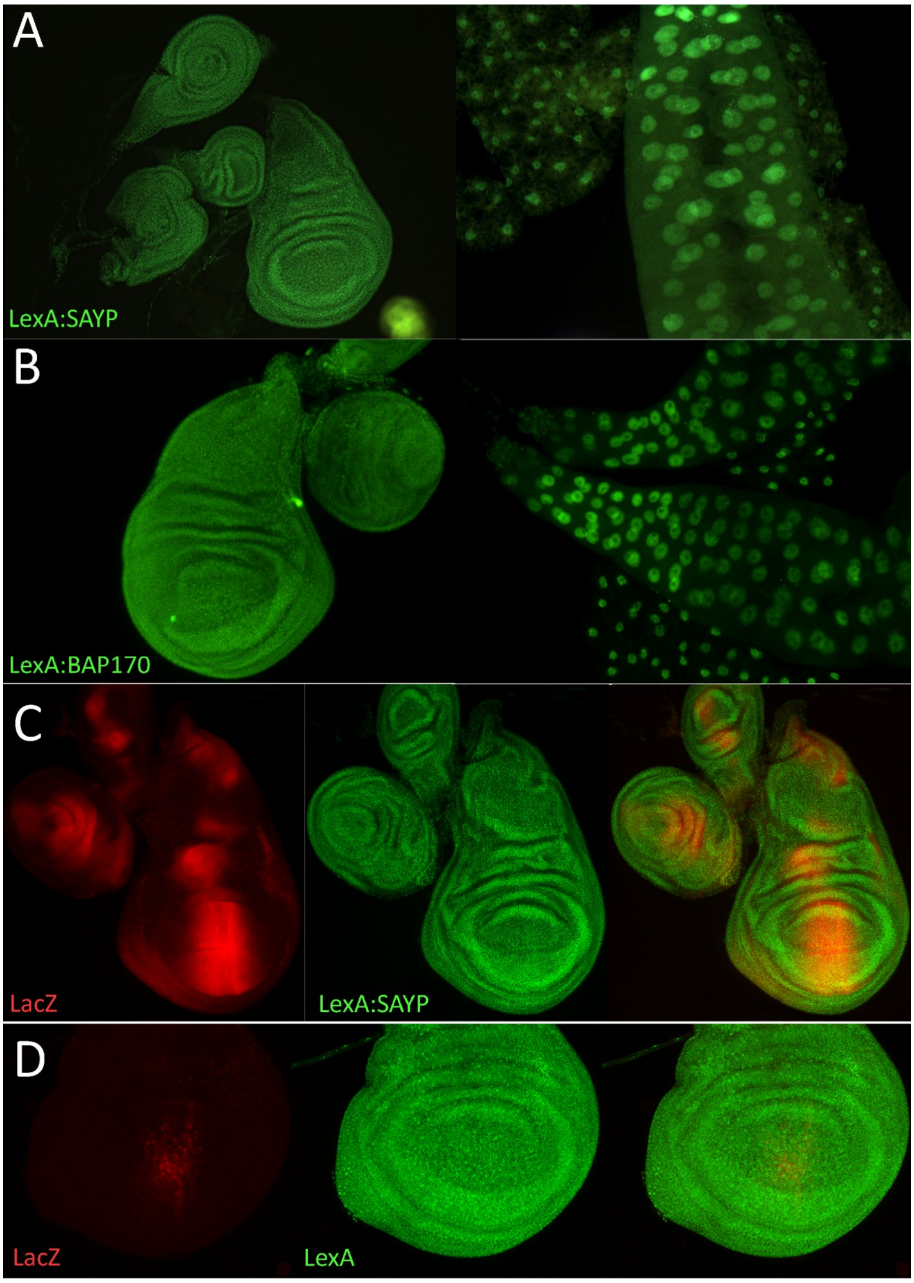


**Figure S1. Ubiquitous expression of LexA:SAYP and LexA:BAP170.**

(A-B) Immunofluorescence imaging of LexA:SAYP and LexA:BAP170 show their ubiquitous expression and nuclear localization in larval tissues (wing imaginal discs are on the left and salivary glands, on the right) of the transgenic stocks carrying P*tub-LexA:SAYP* (A) or P*BAP170-LexA:BAP170*

(B). (C) Double immunolocalization of beta-gal (red) and LexA:SAYP (green) in imaginal discs of transgenic *LexAop-LacZDad* , P*BAP170-LexA:SAYP* larvae shows that, despite the ubiquitous LexA:SAYP expression in wing discs, the *LacZ* reporter is expressed according to the *Dad*-like pattern in a central, vertical stripe of the discs. (D) The LexA protein ubiquitously expressed from the *Ptub-LexA* transgene cannot alone induce expression of the *LexAop-LacZDad* transgene.

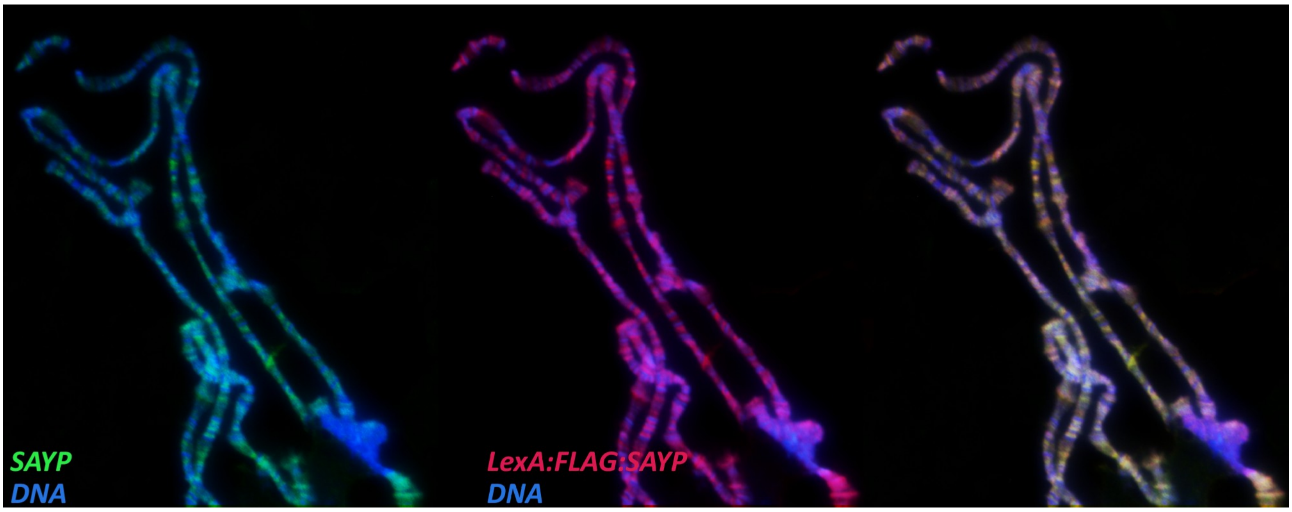


**Figure S2. Colocalization of SAYP and LexA:SAYP on polytene chromosomes.**

The endogenous SAYP (anti-SAYP antibody, left panel, green) and LexA:3xFLAG:SAYP (anti-FLAG antibodies, central panel, red) localize at the same discrete polytenic bands (merge on the right). DAPI staining (blue) marks DNA.

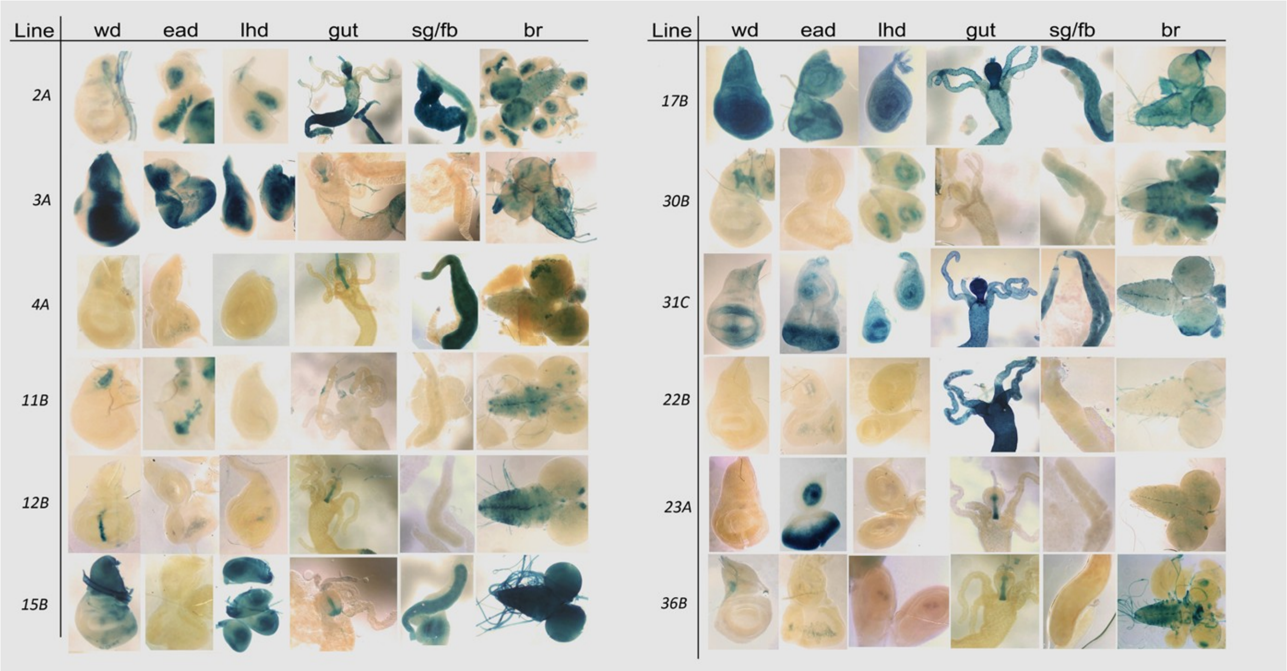


***Figure S3.*** **Beta-gal activity induced in larval tissues by *Ptub-LexA-BAP170/SAYP* in twelve out of twenty lines transgenic for the *LexAop-LacZ* reporter.**

Wd, wing imaginal discs; ead, eye-antennal imaginal discs; lhd, leg-haltere discs; g, guts; sg/fb, salivary glands/fat bodies; br, larval brain.

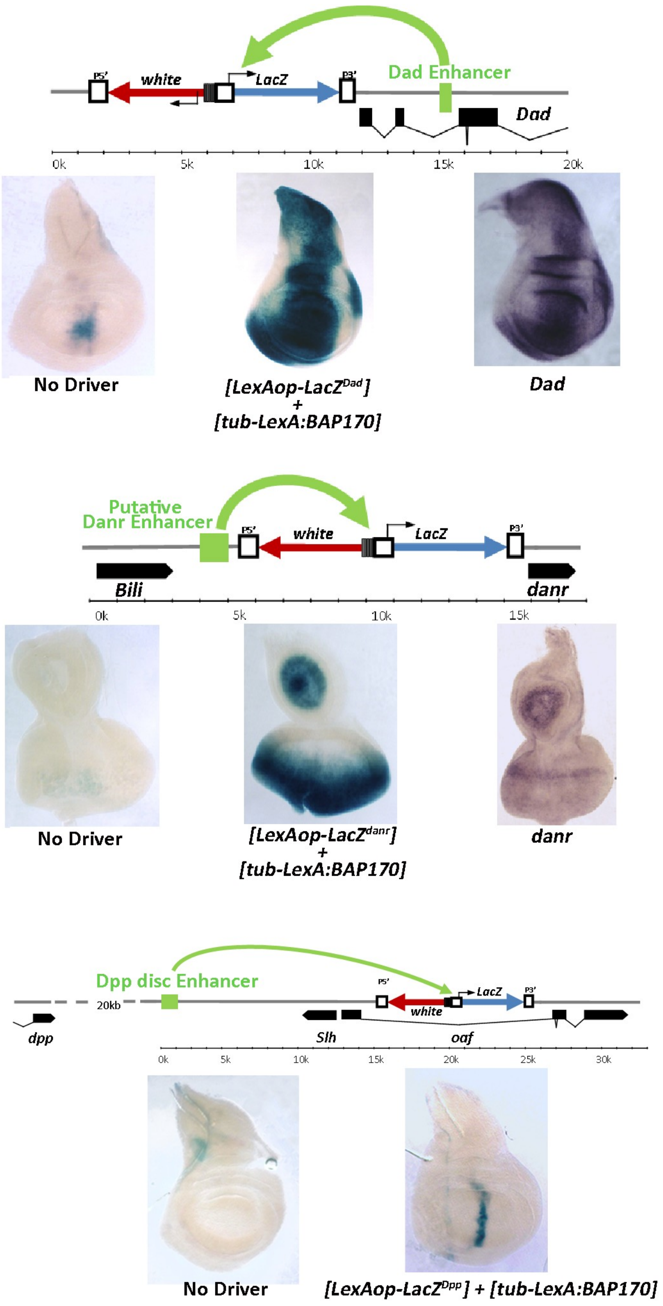


**Figure S4. Enhancer capture by tBAP170 at the *LexAop-LacZDad, LexAop-LacZdanr, and LexAop-LacZDpp* transgenes.**

Schematic drawing of the positions of the *LexAop-LacZ* responders, the enhancers, and flanking genes in the *Dad*, *danr,* and *dpp* genomic regions (top of each panel). Beta-gal activity is detectable in transgenic larvae carrying the specific *LexAop-LacZ* responder alone (left of each panel) or in combination with P*tub*-LexA:BAP170 (center in panels 1 and 2 and right in panel 3). Expression of

*Dad* and *danr* genes, by in situ hybridization, is reported as a control (right in panels 1 and 2); *dpp* expression is not shown.

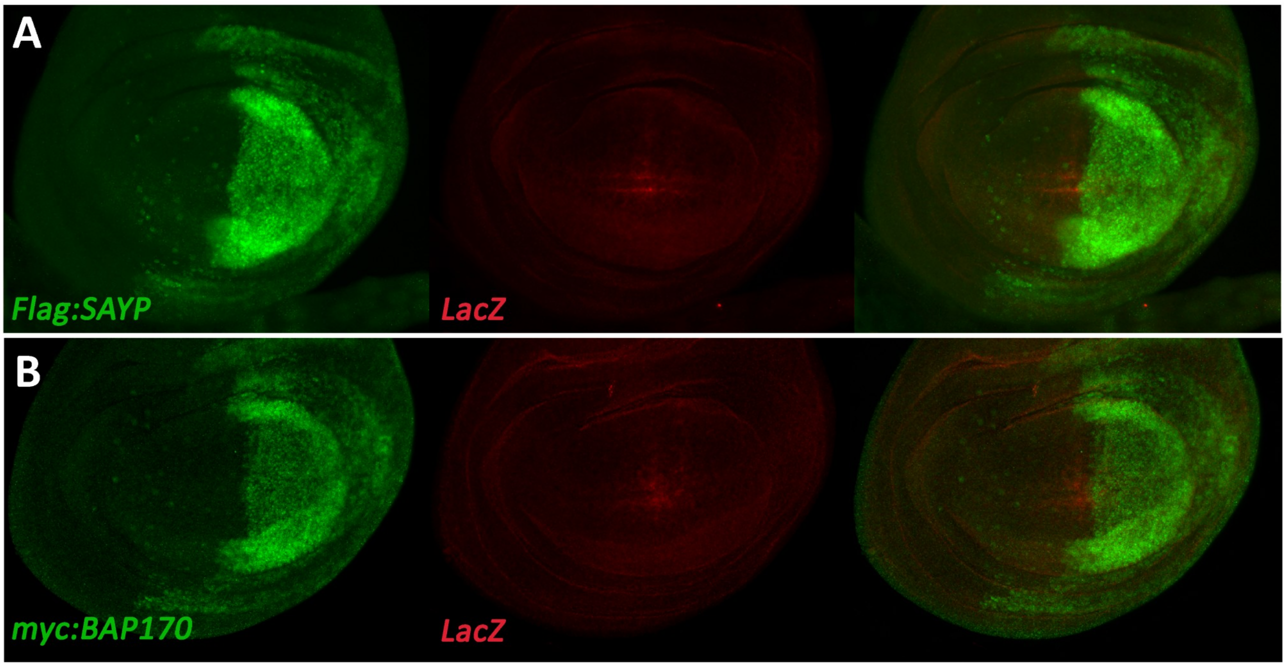


***Figure S5. Wild-type SAYP or BAP170 overexpression cannot induce enhancer-dependent* LexAop-LacZ expression.**

Immuno-fluorescence imaging of wild type SAYP (A) or BAP170 (B) tagged with FLAG or myc epitopes, respectively, expressed from the UAS transgenes under the en-GAL4 driver in the wing discs of *LexAop-LacZDad* transgenic larvae. Expression of the LacZ reporter is not activated by overexpression of SAYP or BAP170 in posterior cells, but remains at its very weak background level. The genotypes are: *UAS-3xFLAG:SAYP/en-GAL4; LexAop-LacZDad* (A), *UAS-myc:SAYP/en-GAL4; LexAop-LacZDad* (B).

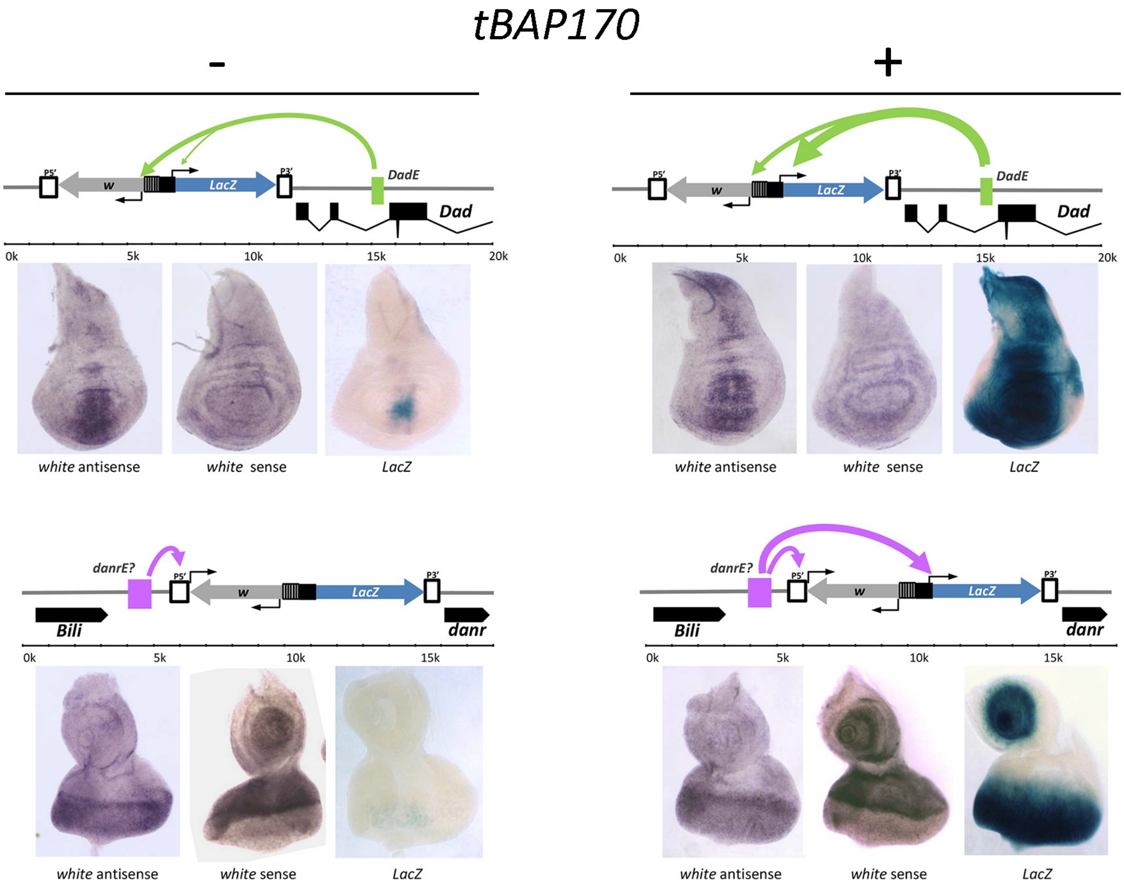


**Figure S6. tBAP170 provides the *hsp70* core promoter with the same tethering activity that is naturally displayed by the white and P-element promoters.**

*The hsp70* core promoter, which lacks a tethering element, is not activated by distant enhancers in the absence of tBAP170 in larvae transgenic for the *LexAop-LacZDad* and *LexAop-LacZdanr* reporters (left panels). Conversely, the *Dad* and *danr* enhancers can spontaneously activate the mini-*white* (top) and P-transposase promoters (bottom). With the same responders, tBAP170 makes the *hsp70* core promoter responsive to both enhancers (right panels). Activities of the mini-*white* and P-transposase promoters were detected by in situ hybridization with an antisense (left of each panel) or a sense probe (center of each panel) to the *white* gene, respectively. Activity of the *hsp70* promoter was monitored by X-gal staining (right of each panel).

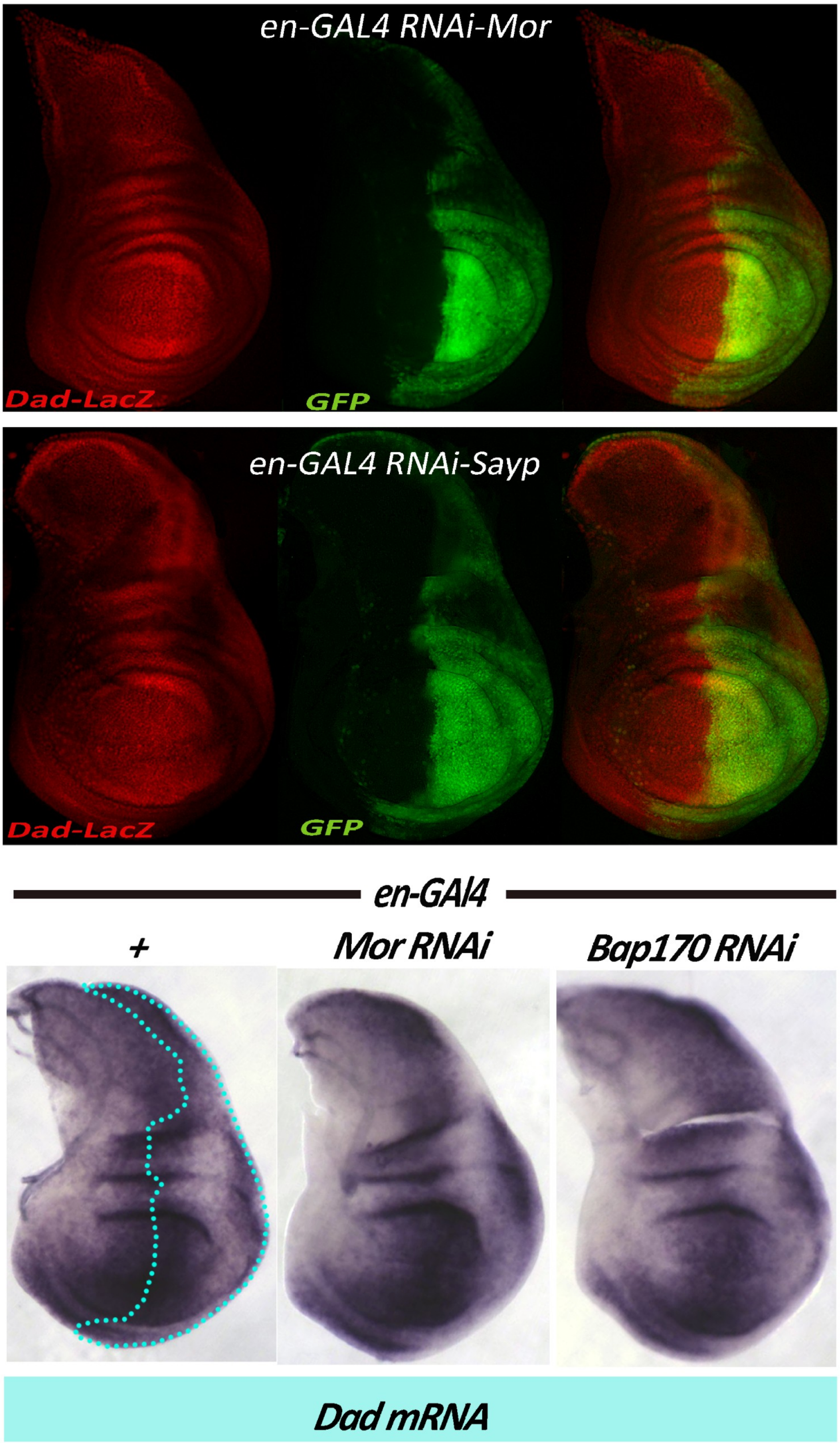


***Figure S7. Dad* expression is independent of PBAP.**

*Dad* enhancer activity is not affected by depletion of MOR (top) or SAYP (middle). *Dad* transcription (bottom panel) is independent of MOR or BAP170 as demonstrated by in situ hybridization with a *Dad* probe. *Dad* enhancer activity was detected using the enhancer trap line *P{lacW}Dadj1E4*. Genotypes: (top) *en-Gal4,UAS-GFP; P{lacW}Dadj1E4/UAS-RNAi-Mor[Vdrc6969]*; (middle) *en-Gal4,UAS-GFP/UAS-RNAi-SAYP [Vdrc105946], P{lacW}Dadj1E4*; (bottom) wild type (left); *en-Gal4; UAS-RNAi-Mor [Vdrc6969]* (center); *en-Gal4, UAS-RNAi-BAP170 [Vdrc34582]* (right).

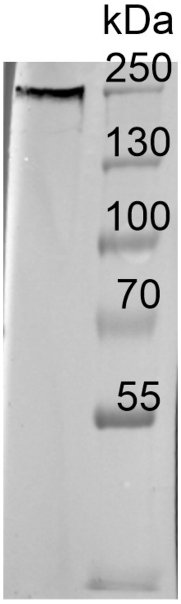


**Figure S8. Western blot analysis of a nuclear embryonic extract with a-BRM antibody.**

| | Affected Gene | RNAi Line | Effect on LexA:Sayp, LexAop-LacZ[Dad] expression | Effect on LexA:Bap170, LexAop-LacZ[Dad] expression | Effect on Dad expression | Phenotype with engrailed-GAL4 (this work) | Phenotype with tubulin-GAL4 (this work) | Phenotypes described with other Gal4 lines |
|---|---|---|---|---|---|---|---|---|
| **PBAP COMPLEX** | *mor* | *VDRC6969* | ↓ | ↑ (Discussed in the text) | N.E. | Prepupal lethal | larval/prepupal lethal | pnr-GAL4[a]<br>Bx-MS1096-GAL4[b]<br>salm-GAL4[c]<br>*C564-Gal4 and Hml-Gal4*[e]<br><br>http://flybase.org/reports/FBal0210097.html |
| | *bap170* | *VDRC34582* | ↓ | ↓ | N.E. | Adult wing extraveins | prepupal lethal | pnr-GAL4[a]<br>salm-GAL4[c]<br><br>http://flybase.org/reports/FBal0198822.html |
| | *brm* | *VDRC37721* | ↓ | N.E. | N.E. | Adult wing defects | larval/prepupal lethal | pnr-GAL4[a]<br>salm-GAL4[c]<br>insc-GAL4[d]<br><br>http://flybase.org/reports/FBal0209177.html |
| | *sayp* | *VDRC105946* | ↓ | N.E. | N.E. | Adult wing defects | larval/prepupal lethal | - |
| | *polybromo* | *VDRC108618* | N.E. | N.E. | N.E. | wt | N.D. | *C564-Gal4 and Hml-Gal4*[e]<br>http://flybase.org/reports/FBal0231490.html |


(a) Mummery-Widmer, J.L., Yamazaki, M., Stoeger, T., Novatchkova, M., Bhalerao, S., Chen, D., Dietzl, G., Dickson, B.J., Knoblich, J.A. (2009).Genome-wide analysis of Notch signalling in Drosophila by transgenic RNAi. Nature 458(7241): 987—992

(b) Dietzl, G., Chen, D., Schnorrer, F., Su, K.C., Barinova, Y., Fellner, M., Gasser, B., Kinsey, K., Oppel, S., Scheiblauer, S., Couto, A., Marra, V., Keleman, K., Dickson, B.J. (2007). A genome-wide transgenic RNAi library for conditional gene inactivation in Drosophila. Nature 448(7150): 151—156

(c) Terriente-Félix, A., de Celis, J.F. (2009). Osa, a subunit of the BAP chromatin-remodelling complex, participates in the regulation of gene expression in response to EGFR signalling in the Drosophila wing. Dev. Biol. 329(2): 350--361.

(d) Neumüller, R.A., Richter, C., Fischer, A., Novatchkova, M., Neumüller, K.G., Knoblich, J.A. (2011). Genome-Wide Analysis of Self-Renewal in Drosophila Neural Stem Cells by Transgenic RNAi. Cell Stem Cell 8(5): 580—593

(e) Bonnay, F., Nguyen, X.H., Cohen-Berros, E., Troxler, L., Batsche, E., Camonis, J., Takeuchi, O., Reichhart, J.M., Matt, N. (2014). Akirin specifies NF-κB selectivity of Drosophila innate immune response via chromatin remodeling. EMBO J. 33(20): 2349--2362.


**Supplementary Table 1**

UAS-RNAi lines used in this work. Effects on LexA-SAYP/LexA-BAP170 mediated activation of the *LexAop-LacZDad* responder are indicated (↓, downregulated; ↑, upregulated; NE, no effect; ND, not determined). All lines except UAS-RNAi-Polybromo cause an evident phenotype upon activation with en-Gal4 (from larval/prepupal lethality to defects in adult wing morphology, columns 7 and 8). Phenotypes induced with other GAL4 lines are described in the last column, with references.

### Supplementary materials

**Primers for PCR amplification and cloning of the *Dad* and *dpp* enhancers**

JFL-attB [DadEnh-lexAop-hsp70-LacZ], 5’-cgAAGCTTTCGAGAGCGCCTTCAATTCTAATTC and 5’-cgAAGCTTTTCACCAACGGACAGCCGACGC

JFL-attB [lexAop-hsp70-LacZ-DadEnh], 5’-cgGAATTCTCGAGAGCGCCTTCAATTCTAATTC and 5’- cgGAATTCTTCACCAACGGACAGCCGACGC

JFL-attB [dppEnh-lexAop-hsp70-LacZ], 5’- cgAAGCTTTTCCACTCACCTTGTCAGCCAG and 5’- cgAAGCTTATTCCAGTGCTGGGAACGTG

JFL-attB [lexAop-hsp70-LacZ-dppEnh], 5’- cgGAATTCTTCCACTCACCTTGTCAGCCAG and 5’- cgGAATTCATTCCAGTGCTGGGAACGTG

**Primers used in ChIP**

point 1 (*hsp70* reporter promoter), 5’-TAGCGCTAGCGACGTCGAG and 5’-GCTTAGCGACGTGTTCACTTTG

point 2, 5’-TGTGCGATATATGGCTTTTGAAGTG and 5’-TGTAATCCAGTGAACAATAGAGAGAGCTG

point 3 (*Dad* enhancer), 5’-AGGTGAGCGTGTGTTGGTGTG and 5’-TCATACATACAGAATGCTTGCGTGC

point 4, 5’-TTGCCCATACCGAAAAATCCTG and 5’-TCAGCTACAGTGAAGCGTCTCAAAC

rDNA, 5’-AATTCAGAACTGGCACGGACTTGG and 5’-AGAGCACTGGGCAGAAATCACATTG
